# Switchable microscale stress response of actin-vimentin composites emerges from scale-dependent interactions

**DOI:** 10.1101/2024.06.07.597906

**Authors:** Julie Pinchiaroli, Renita Saldanha, Alison E Patteson, Rae M. Robertson-Anderson, Bekele J. Gurmessa

**Affiliations:** Department of Physics and Astronomy, Bucknell University, Lewisburg, PA, 17837, USA; Department of Physics and BioInspired Institute, Syracuse University, Syracuse, NY 13210, USA; Department of Physics and Biophysics, University of San Diego, San Diego, CA 92110, USA

## Abstract

The mechanical properties of the mammalian cell regulate many cellular functions and are largely dictated by the cytoskeleton, a composite network of protein filaments, including actin, microtubules, and intermediate filaments. Interactions between these distinct filaments give rise to emergent mechanical properties that are difficult to generate synthetically, and recent studies have made great strides in advancing our understanding of the mechanical interplay between actin and microtubule filaments. While intermediate filaments play critical roles in the stress response of cells, their effect on the rheological properties of the composite cytoskeleton remains poorly understood. Here, we use optical tweezers microrheology to measure the linear viscoelastic properties and nonlinear stress response of composites of actin and vimentin with varying molar ratios of actin to vimentin. We reveal a surprising, nearly opposite effect of actin-vimentin network mechanics compared to single-component networks in the linear versus nonlinear regimes. Namely, the linear elastic plateau modulus and zero-shear viscosity are markedly reduced in composites compared to single-component networks of actin or vimentin, whereas the initial response force and stiffness are maximized in composites versus single-component networks in the nonlinear regime. While these emergent trends are indicative of distinct interactions between actin and vimentin, nonlinear stiffening and longtime stress response appear to both be dictated primarily by actin, at odds with previous bulk rheology studies. We demonstrate that these complex, scale-dependent effects arise from the varied contributions of network density, filament stiffness, non-specific interactions, and poroelasticity to the mechanical response at different spatiotemporal scales. Cells may harness this complex behavior to facilitate distinct stress responses at different scales and in response to different stimuli to allow for their hallmark multifunctionality.

## I. INTRODUCTION

The cytoskeleton of mammalian cells is comprised of three distinct filamentous proteins: actin, microtubules, and intermediate filaments (IFs). These filaments work together to provide cells with mechanical integrity while also regulating myriad mechanical processes such as cell adhesion, cell division, nuclear positioning, and migration [1–7]. Each of these filaments has distinct stiffness, size, and stability that dictate their different functions. More recently, it has become increasingly appreciated that interactions and crosstalk between the different filaments are equally critical to many mechanical processes [8–13]. Similarly, in materials applications, composites of polymers of varying stiffnesses are often employed to evade the limitations of single-component networks, allowing for emergent mechanical properties such as increased resilience and tunability [9, 12, 14, 15].

Semiflexible actin filaments and rigid microtubules are highly preserved in cells and well characterized, and their mechanical properties and cellular functions of their networks have been extensively investigated both *in vivo* and *in vitro* [4, 8, 11]. Conversely, intermediate filaments constitute an entire class of 50 different proteins, with each type having different structural and mechanical properties [16–18]. Even a single IF type, such as vimentin, which is expressed in mesenchymal cells and plays key roles in cell adhesion, migration, and mechanosensing [19–23], can adopt a wide range of lengths and conformations and may non-specifically crosslink with neighboring IFs or other proteins [8, 18, 24, 25]. This diversity allows IFs to contribute to many different cellular functions and has in part obscured how their mechanical properties enable these different functionality in cells [26]. Nevertheless, previous studies have suggested that IFs are essential to the hallmark stress stiffening and recovery behavior of cells [7, 17, 26–28]. How these distinct mechanical features rely on or are impacted by the presence of the other cytoskeletal filaments remains poorly understood.

Here, we address this problem by determining the linear and nonlinear microrheological properties of composites of actin filaments and vimentin IFs with varying molar fractions of each component, revealing intriguing emergent mechanical properties that are highly dependent on the spatiotemporal strain characteristics. Semiflexible actin filaments, which play pivotal roles in cellular functions, ranging from tensile strength and apoptosis to motility and replication, have persistence lengths of *l*_*p*_ ≃ 10 *μ*m, comparable to their typical contour lengths *L* ≈ 5 − 20 *μ*m [1–4, 29–31]. In vitro, entangled networks of actin filaments exhibit linear viscoelastic properties that can be described by extensions of the reptation model to semiflexible polymers, with a low-frequency elastic plateau modulus that scales as *G*^0^ ∼ *c*^1.4^ [32, 33]. The nonlinear response of entangled actin networks is more complex, with networks exhibiting varying degrees of stress stiffening and softening, as well as different relaxation mechanisms and timescales, depending on the spatiotemporal scale of the strain and the actin concentration [34–40].

Vimentin forms a meshwork that envelops actin bundles and stress fibers in cells [5, 41]. Vimentin IFs have similar contour lengths as actin filaments but are much more flexible, with *l*_*p*_ ≃ 2 *μ*m. They have been shown to be uniquely extensible, a feature which has been implicated in their excepted role in stress absorption, deformability, and stiffening of cells [25, 42–44]. Reconstituted vimentin IF networks exhibit minimal stress relaxation in the linear regime, with linear viscoelastic moduli that display very weak scaling with frequency. The concentration dependence of *G*^0^ has also been shown to be weaker than that for actin networks, with *G*^0^ ∼ *c*^0.47^ reported for bulk rheology measurements [20]. Above a critical strain, vimentin networks exhibit stress stiffening that can be described as *K* ∼ *σ*^3*/*2^ where *K* is the nonlinear modulus and *σ* is stress [45]. Moreover, the degree of stiffening and critical stiffening strain are dependent on the concentration of vimentin and divalent salts [25].

Despite their significance to cellular processes and materials applications, the mechanical properties of composites of actin and vimentin are not fully understood. Bulk rheological measurements have reported nonmonotonic dependence of the elastic modulus *G*^′^ on the molar fraction of actin in the composite *ϕ*_*A*_, reaching a pronounced peak at *ϕ*_*A*_ = 0.25, which was attributed to non-specific crosslinking between actin and the tail-domain of vimentin [18]. Furthermore, bulk measurements on composites of actin and another IF, keratin, reported similar albeit weaker non-monotonicity [9, 46, 47]. However, similar studies performed on actin-VIF composites at lower protein and salt concentrations reported a monotonic increase in *G*^′^ with *ϕ*_*A*_ [16]. Theoretical models that recapitulated this monotonic trend incorporate non-specific interactions between vimentin IFs but no actin-vimentin interactions [18, 48]. In response to nonlinear bulk strains, the addition of vimentin to crosslinked actin has been reported to either increase or decrease the degree of stress stiffening, dependent on the actin concentration. These findings, captured by theoretical models, were shown to arise from the ability of vimentin to either suppress or enhance actin crosslinking, depending on the actin and crosslinker concentrations. Finally, particletracking microrheology experiments of VIF networks at fixed concentration with varying concentrations of actin added reported scaling 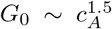, similar but slightly larger than 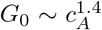 reported for actin networks in the absence of VIFs [26, 49].

These few studies, reporting complex and conflicting results, underscore our limited understanding of the mechanics of actin-IF composites and the need for microscale and mesoscale rheology measurements that are equipped to delineate the various contributions to the varied bulk response features reported.

In this manuscript, we use optical tweezers microrheology to characterize the linear and nonlinear mechanical response of *in vitro* composites of co-entangled actin filaments and vimentin IFs (Fig 1). We quantify the frequency-dependent viscoelastic moduli *G*^′^(*ω*) and *G*^*′′*^(*ω*) from the thermal fluctuations of optically trapped microsphere probes embedded in the composites. We also further characterize the nonlinear response by optically displacing probes through the composites and measuring the forces the composites exert on the probes during and after the strain (Fig 1B-D). We show that the dependence of rheological properties on composite composition (*i*.*e*., fraction of actin *ϕ*_*A*_) is markedly different in the linear and nonlinear regime as well as at different timescales. Namely, composites displayed reduced elastic moduli and zero-shear viscosity compared to single-component networks, while short-time nonlinear stress response and stiffness are enhanced in composites. These nearly opposite emergent behaviors suggest a careful interplay between network connectivity and filament rigidity, as well as the prevalence of easily disrupted IF interactions. Moreover, we demonstrate that the actin fraction dictates the long-time behavior and degree of stress-stiffening, at odds with current thought based on bulk rheology experiments. Our results suggest that cells may utilize this complex behavior to enable various stress responses at different scales and in reaction to various stimuli, allowing for their versatile functionality.

**FIG. 1.**
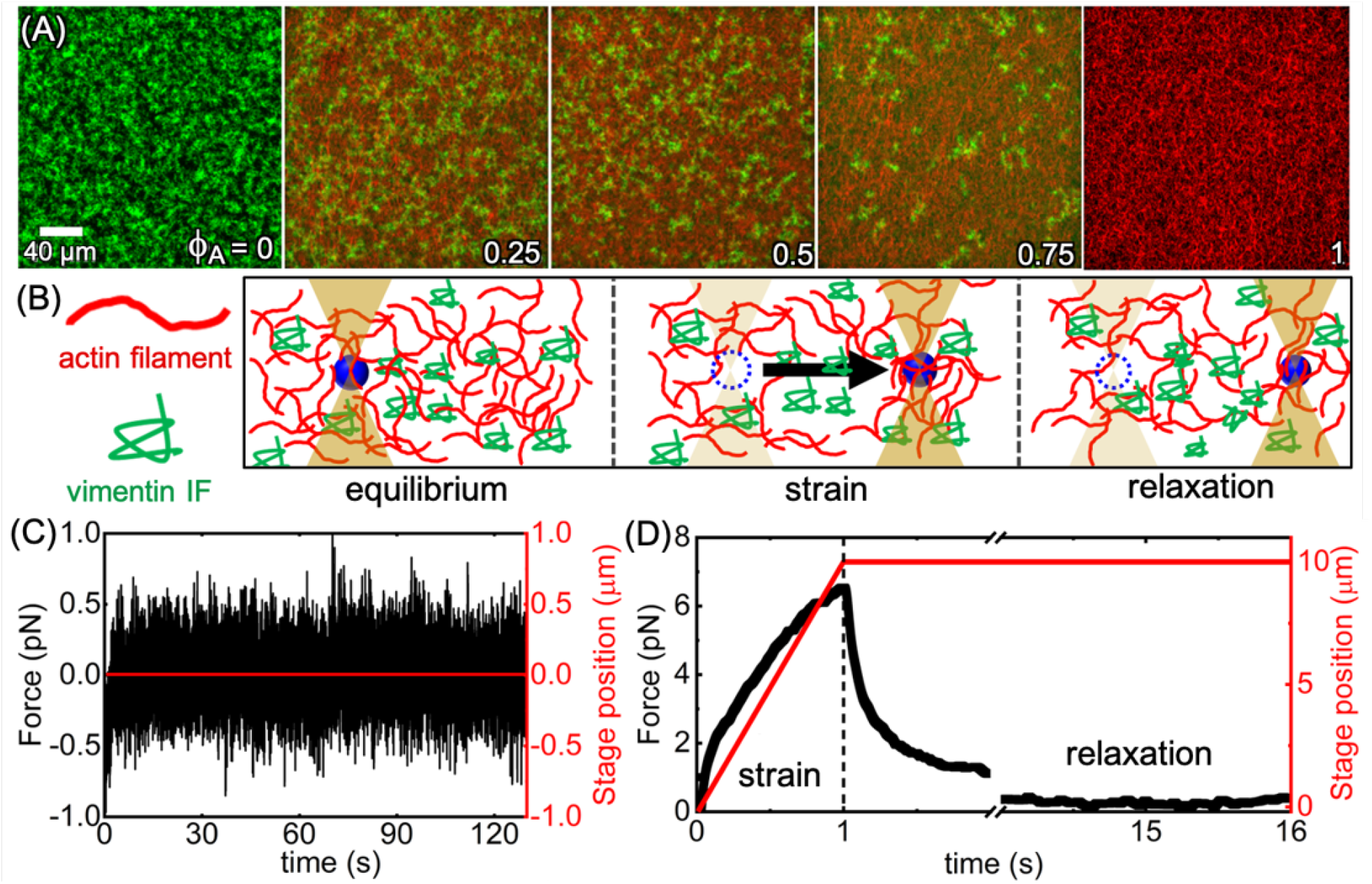
Design and optical tweezers microrheology of entangled composites of actin filaments and vimentin intermediate filaments. (A) Composite networks of actin filaments (red) and vimentin intermediate filaments (green) prepared at fixed total protein concentration *c*_*A*_ + *c*_*V*_ = 11.6 *μ*M with varying molar fractions of actin *ϕ*_*A*_ = *c*_*A*_*/*(*c*_*A*_ + *c*_*V*_) = 0, 0.25. 0.5, 0.75 and 1 (left to right). Each image is a *z*-projection of a stack of 35 512×512 square-pixel images separated in *z* by 0.5 *μ*m, collected with a Leica TCS SP5 laser scanning confocal microscope with 63× 1.4 NA objective. (B-D) Optical tweezers microrheology measurements of actin-vimentin composites. (B) Cartoon depicting a microsphere probe (blue) of diameter *d* = 4.2 *μ*m embedded in an actin-vimentin composite and trapped by a focused laser beam during the three phases of an OTM experiment: (equilibrium) the probe is held fixed, (strain) the probe is displaced 10 *μ*m through the composite at a constant speed of *v* = 10 *μ*m s^−1^ by moving the sample stage relative to the trap, and (relaxation) the trap is held fixed following the strain as the probe relaxes back to the center of the trap. The force exerted on the probe during all phases is collected at 20 kHz. (C) The equilibrium phase is used to determine linear viscoelastic properties. For linear OTM measurements, the stage position (red) is held fixed, and the thermal force fluctuations (black) are measured for 130 s to extract frequency-dependent linear viscoelastic moduli *G*^′^(*ω*) and *G*^*′′*^(*ω*). (D) For nonlinear OTM measurements, the force exerted on the probe (black) is measured during the strain and relaxation phases, during which the stage position (red) increases and is then held constant. Sample data shown in C,D are for the *ϕ*_*A*_ = 0.75 composite.

## II. RESULTS

### Design of interpenetrating networks of actin and vimentin that mimic in vivo structures

To decipher the distinct contributions from actin and vimentin and the emergent interactions between the two filament types, we prepare composites with fixed total protein concentration of *c* = *c*_*V*_ + *c*_*A*_ = 11.6 *μ*M and varying molar fractions of actin *ϕ*_*A*_ = *c*_*A*_*/c* and vimentin *ϕ*_*V*_ = 1 − *ϕ*_*A*_ (Fig 1). Two-color confocal imaging experiments show that single-component networks (actin or vimentin only) and composites at all actin fractions *ϕ*_*A*_ comprise overlapping actin and vimentin filaments that are homogenously distributed across mesoscopic scales (∼200 *μ*m).

We first aim to confirm that our composite preparation methods, which result in homogeneous interpenetrating networks of actin filaments and vimentin intermediate filaments, are physiologically relevant [5]. To demonstrate this relevance, we image actin filaments and vimentin IFs in mouse embryonic fibroblasts (mEF), as shown by the immunofluorescence images of an mEF, stained with spectrally-distinct fluorescent phalloidin and antivimentin antibodies that bind to actin and vimentin, respectively (Fig 2A). While cellular actin and vimentin networks both appear to be heterogeneously distributed, there are distinct regions that comprise interpenetrating networks of actin and vimentin, qualitatively similar to those we observe *in vitro* (Fig 1A).

**FIG. 2.**
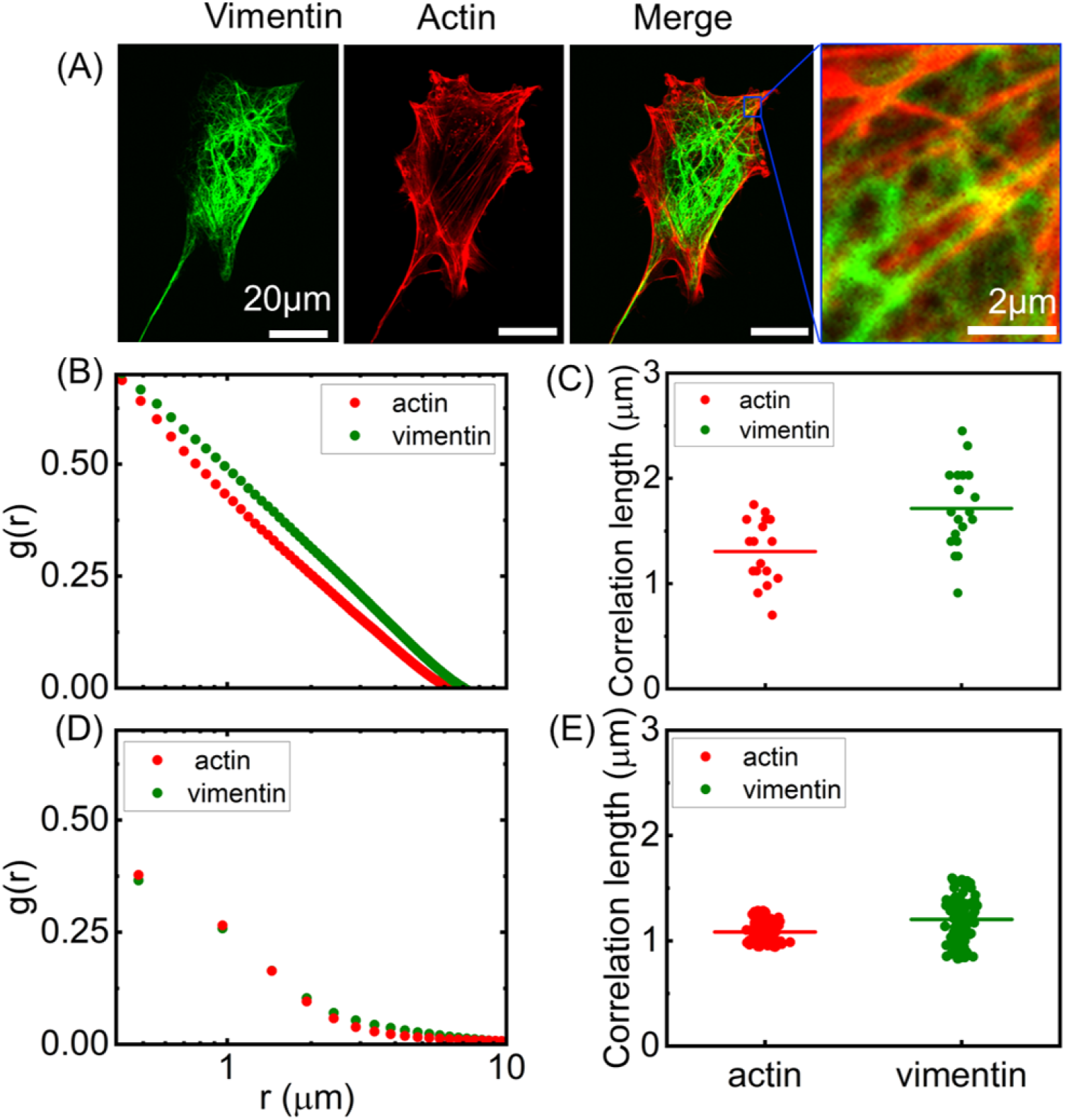
Interpenetrating networks of actin and vimentin in cells have similar structural features as in vitro composites. (A) Immunofluorescence images of mouse embryonic fibroblasts (mEF) labeled for vimentin (left, green) and actin (red, middle left), with the two channels merged (middle right) and zoomed-in (right), respectively, (B-E) Spatial image autocorrelation analysis is performed on images of composites in mEF cells (B,C) and compared to in vitro composite images (D,E). (B) Spatial image autocorrelation function *g*(*r*) computed for the separate actin (red) and vimentin (green) channels of the in vivo fluorescence images shows a similar structural correlation for actin and vimentin that decays with increasing distance *r*. The characteristic distance *r* associated with the decay is a measure of the structural correlation length *ξ*_*g*_ for each filament network, shown in (C) for all images (circles) with the horizontal bars denoting the average *ξ*_*g,A*_ and *ξ*_*g,V*_. The data shows that both the average and spread in *ξ*_*g*_ values are modestly higher for vimentin compared to actin. (D,E) The same metrics extracted from cell images performed on fluorescence confocal images of *in vitro* composites (Fig 1A) show (D) *g*(*r*) curves and (E) *ξ*_*g*_ values for actin (red) and vimentin (green) have similar features to those measured in cells, with vimentin exhibiting slightly higher average *g*(*r*) and *ξ*_*g*_ values and greater spread across different images. Data shown in (D,E) is an average over all data for *ϕ*_*A*_= 0.25, 0.5, 0.75 composites.

To quantify the structural characteristics of these composite regions, we perform spatial image autocorrelation (SIA) analysis (see Methods) on the actin and vimentin images to determine a distinct intensity autocorrelation function *g*(*r*) for each network. The *g*(*r*) data quantifies the extent to which two pixels separated by a distance *r* are correlated and generally displays exponential decay with *r*. The critical lengthscale associated with the exponential decay can be understood as the characteristic structural correlation lengthscale *ξ*_*g*_, which, for a homogoneous network, can serve as a proxy for mesh size (Fig 2). As shown in Fig 2B, average *g*(*r*) curves are similar for both filament types, but the vimentin structure appears to decay more weakly with increasing *r*. The resulting correlation lengths, determined from fits of the individual *g*(*r*) curves that constitute the averages shown in Fig 2C, show that vimentin has a modestly higher correlation length of *ξ*_*g,V*_ = 1.75 *±* 0.09 *μ*m compared to *ξ*_*g,A*_ = 1.25 *±* 0.07 *μ*m (p *<* 0.01).

We perform the same SIA analysis on our *in vitro* composite images, finding qualitatively similar *g*(*r*) features as the *in vivo* measurements, with comparable but slightly lower correlation lengths of *ξ*_*g,A*_ = 1.1 *±* 0.01 *μ*m and *ξ*_*g,V*_ = 1.2 *±* 0.02 *μ*m for actin and vimentin, respectively. Moreover, in both *in vitro* and *in vivo* measurements, the spread in *ξ*_*g,V*_ values is larger than that for actin, indicative of more heterogenous distribution of vimentin IFs compared to actin in composites. These results demonstrate that our designed in vitro composites are structurally similar to *in vivo* networks, suggesting that the microscale mechanical properties we investigate may provide valuable new insights into cellular mechanics.

### Actin-vimentin composites exhibit emergent non-monotonic viscoelastic properties depending on composite composition

We characterize the linear viscoelastic properties of the composites shown in Fig 1A by analyzing the thermal fluctuations of optically trapped microspheres embedded in composites, as depicted in Fig 1B,C. Using the generalized Stokes-Einstein relationship (GSER), we determine the frequency-dependent linear elastic and viscous moduli, *G*^′^(*ω*) and *G*^′′^(*ω*) for networks of either actin (*ϕ*_*A*_ = 1) or vimentin (*ϕ*_*V*_ = 0) (Fig 3A) compared to composites (*ϕ*_*A*_ = 0.25, 0.5, 0.75) (Fig 3B). We first observe that both the magnitude and frequency dependence of *G*^′^(*ω*) and *G*^*′′*^(*ω*) are surprisingly similar for actin and vimentin networks despite the order of magnitude difference in their persistence lengths. Moreover, Fig 2 suggests that the mesh size of the vimentin network is slightly larger, which we may expect to result in lower *G*^′^(*ω*). Likewise, using predicted expressions for the mesh size of actin and vimentin networks with mass concentration *c*_*A*_ and *c*_*V*_, 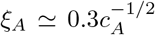 and 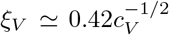, where *c*_*A*_ and *c*_*V*_ are in units of mg mL^−1^, and the corresponding mesh sizes are in units of *μ*m (see Methods, SI Fig S3), we compute *ξ*_*A*_ ≃ 0.42 *μ*m which is smaller than *ξ*_*V*_ ≃ 0.52 *μ*m [16, 50, 51]. We also calculate the crossover times, (*τ*_*c*1_ & *τ*_*c*2_) (Fig 3E), the frequency dependent shear viscosity, *η*(*ω*), and the loss tangent, tan *δ* (Methods, SI Fig S1(B,C)).

**FIG. 3.**
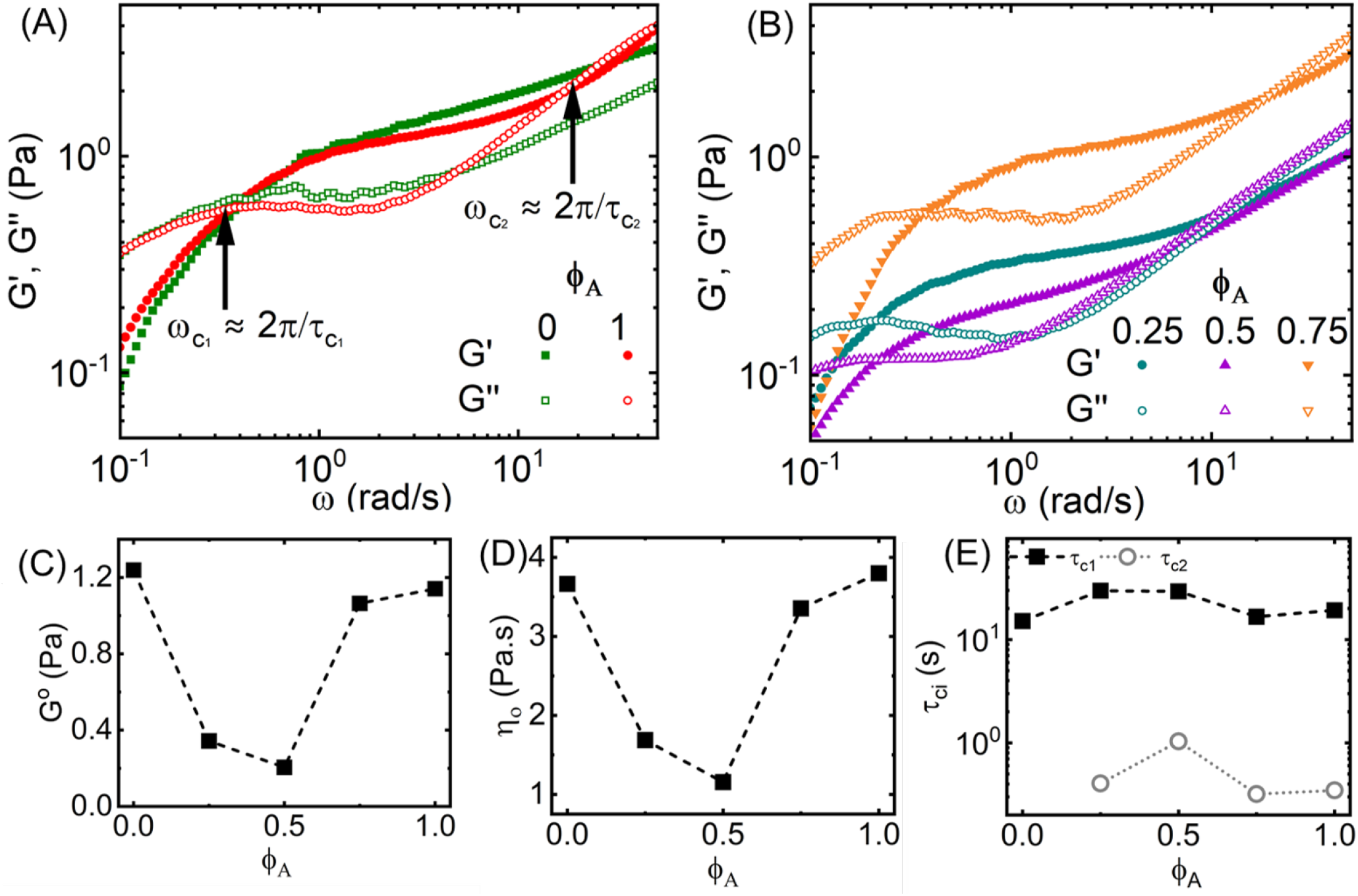
Linear microrheological properties of actin-vimentin composites exhibit non-monotonic dependence on composite composition. (A,B) Linear frequency-dependent elastic (filled symbols) and viscous (open symbols) moduli, *G*^′^(*ω*) and *G*^′′^(*ω*), for (A) single-component networks of actin filaments (*ϕ*_*A*_ = 1, red) and vimentin IFs (*ϕ*_*A*_ = 0, green) compared to (B) actin-vimentin composites with *ϕ*_*A*_ = 0.25 (dark cyan), 0.5 (purple) and 0.75 (orange). (C) Plateau modulus *G*^0^ and (D) zero-shear viscosity *η*_0_ display similar non-monotonic dependence on molar actin fraction *ϕ*_*A*_, reaching minima at *ϕ*_*A*_ = 0.5. (E) The relaxation times 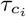 determined from the frequencies 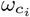 at which *G*^′^(*ω*) = *G*^′′^(*ω*) via the relation 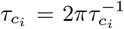. Example slow and fast crossover frequencies, 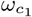 and 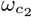, which correspond to the long and short relaxation times 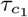 and 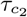 are denoted in (A).

The similarity between the two single-composite actin and vimentin networks is strongest at lower frequencies, where both networks exhibit terminal flow behavior, with *G*^′′^(*ω*) *> G*^′^(*ω*), followed by a crossover to an elasticdominated regime at 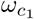, where *G*^′^(*ω*) becomes greater than *G*^′′^(*ω*). This crossover frequency is a measure of the longest relaxation time of the network 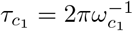 which is the disengagement time *τ*_*D*_ for entangled polymers. Namely, *τ*_*D*_ is the time over which an entangled polymer is able to diffuse out of its deformed entanglement tube and into a pristine tube [35, 37, 52]. We measure similar values of 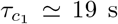 and ∼15 s for actin and vimentin networks, respectively. We also find comparable values for the elastic plateau modulus *G*^0^ for the two networks, 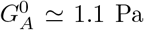 and 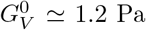 which we determine by evaluating *G*^′^ at the frequency at which the loss tangent is minimum (Fig 3C, SI Fig S1B) [53, 54].

Deviations between the two networks emerge at higher frequencies, *ω* ≳ 3 rad s^−1^, where the actin network exhibits a second crossover to a viscous-dominated regime at 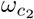, a signature of entangled polymers at timescales below the entanglement time *τ*_*e*_, where mechanics are dominated by single chains that do not feel the constraints of their neighbors [52]. The lack of a crossover for vimentin networks is suggestive of tighter constraints and extended timescales over which interactions with neighboring chains dominate the mechanics, ostensibly at odds with the larger mesh size of vimentin compared to actin. Turning our attention to the composites, we find similar frequency dependence as single-component networks for all *ϕ*_*A*_, including a low-frequency flow regime, a crossover at 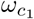, and elastic plateau regime defined by a plateau modulus *G*^0^ (Fig 3B). Similar to actin networks, all composites also exhibit a high-frequency crossover at 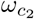 that is absent in the vimentin-only (*ϕ*_*A*_ = 0) network. This result may indicate that the relaxation mechanism related to 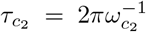 is a signature of the actin in the composites or that the mechanism preventing this crossover in *ϕ*_*A*_ = 0 is disrupted or suppressed.

While the frequency dependence of *G*^′^(*ω*) and *G*^′′^(*ω*) is similar for composites and single-component networks, their magnitudes are universally lower for composites. This effect can be seen clearly in Fig 3C which shows a strong non-monotonic dependence of the plateau modulus *G*^0^ on *ϕ*_*A*_, with *G*^0^ dropping by a factor of ∼6 from *G*^0^ ≃1.2 Pa at *ϕ*_*A*_ = 0 to ∼0.2 Pa at *ϕ*_*A*_ = 0.5 followed by a increase back to ∼1.1 Pa at *ϕ*_*A*_ = 1. We observe a very similar non-monotonic dependence of the zero-shear viscosity *η*_0_, which we approximate as the complex viscosity *η* at the lowest frequency we measure (Fig 3D, SI Fig S1).

Finally, we examine the *ϕ*_*A*_-dependence of the slow and fast relaxation timescales, 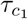 and 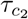 determined from the respective crossover frequencies. We find weaker dependence on *ϕ*_*A*_ compared to the moduli magnitudes (Fig 3C,D), particularly for the slower timescale, which we expect to be related to the disengagement time *τ*_*D*_. The fast timescale 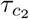 shows a peak at *ϕ*_*A*_ = 0.5, indicative of a longer time required for the onset of entanglement dynamics if we assume 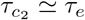, a result that aligns with the corresponding minimum in *G*^0^.

Our collective linear regime results suggest that composites have weaker constraints and are more easily able to dissipate local stresses compared to single component networks, ostensibly at odds with previous bulk rheology measurements [18, 19], which may arise from increased mesh size and/or suppressed non-specific vimentin interactions (i.e., crosslinking). However, the similarity in the longest measured relaxation timescales across the composition space suggests that universal relaxation mechanisms may be at play. We explore these possibilities in later sections.

### Nonlinear straining induces emergent stiffness of actin-vimentin composites that is quenched at large strains

To determine the extent to which the behavior we observe in the linear regime holds when composites are subject to large nonlinear strains that push them out of equilibrium, we perform nonlinear constant rate OTM measurements (Fig 1B,D). In each of these measurements, the same optically trapped probe used for linear OTM is displaced a distance *x*_*max*_ = 10 *μ*m through the composite at constant speed *v* = 10 *μ*m s^−1^, after which the probe is held fixed for 15 s. We measure the force the composite exerts to resist the strain and the subsequent relaxation of the force as the probe is held fixed. In response to the strain, all networks exhibit a sharp rise in force, followed by a more shallow rise with increasing distance *x*, as shown in Figure 4A. For *ϕ*_*A*_ *<* 0.5, the response force reaches a nearly strain-independent plateau, indicating a largely viscous terminal response. However, as *ϕ*_*A*_ increases, the large-strain response force becomes increasingly dependent on *x* (*i*.*e*., the slope of *F* (*x*) increases), a signature of increasing elasticity or stiffness. The magnitude of *F* (*x*) likewise increases with *ϕ*_*A*_, a dependence that we quantify by evaluating the terminal value of the force at the end of the strain *F*_*t*_ = *F* (*x* = 10 *μ*m) as a function of *ϕ*_*A*_ (Fig 4B). We find that *F*_*t*_ follows power-law scaling 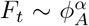 with scaling exponent *α* ≃ 1.4 that aligns with the scaling of the shear modulus with concentration for entangled actin networks [51, 55, 56], suggesting that actin dominates the long-time behavior in the nonlinear regime.

**FIG. 4.**
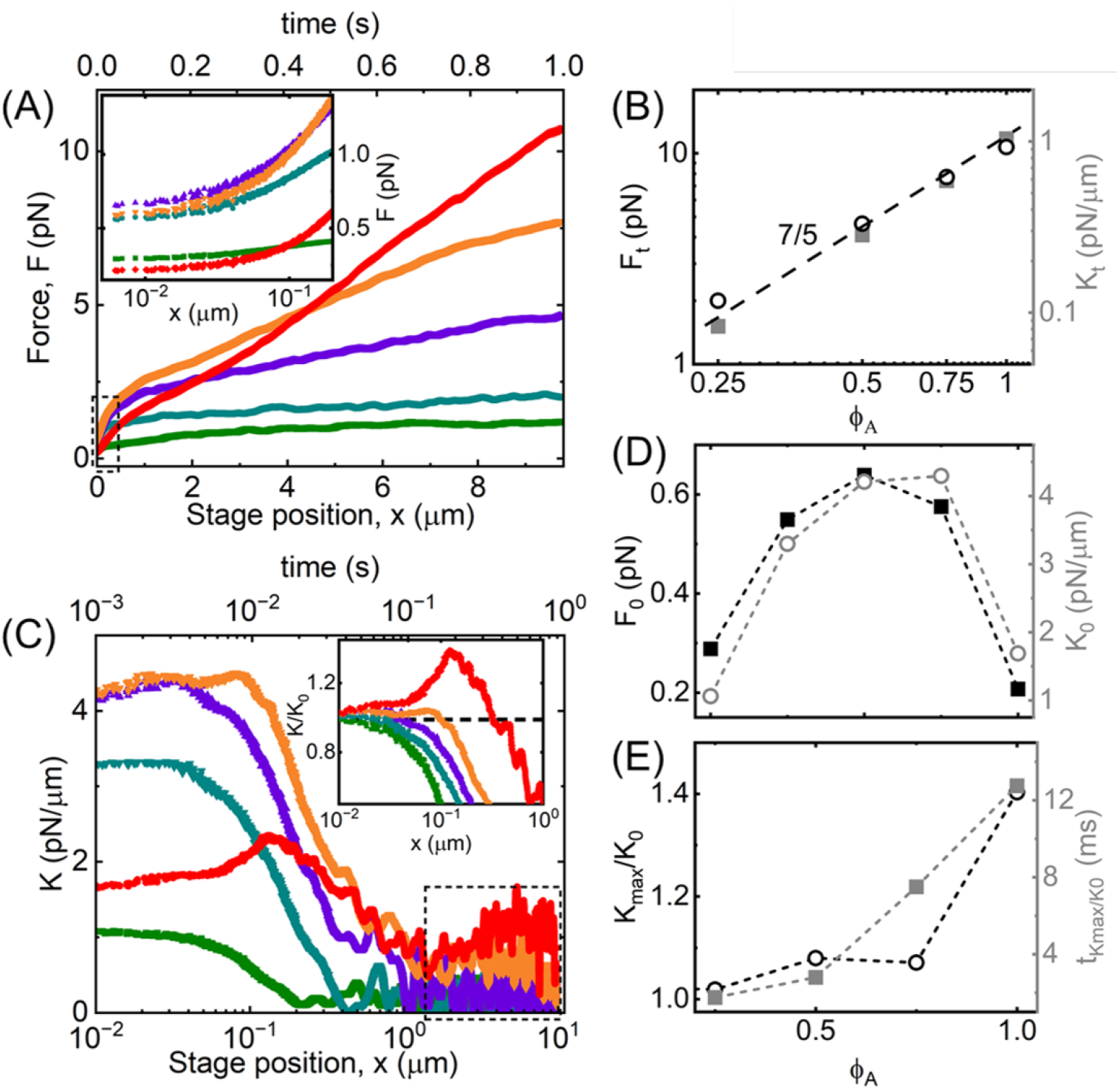
Nonlinear microrheology of actin-vimentin composites manifests emergent stiffness not accessible in the linear regime that is suppressed at large strains. (A) The force exerted by composites with *ϕ*_*A*_ = 0 (green), 0.25 (dark cyan), 0.5 (purple), 0.75 (orange), and 1 (red) to resist the moving probe as it is displaced a distance *x* (bottom axis) over a time *t* = *x/v* (top axis) at constant speed *v* = 10 *μ*m s^−1^. (Inset) Zoom-in of the small-*x* region denoted by dashed box to show the dependence of the initial force *F*_0_ on *ϕ*_*A*_. (B) Terminal force *F*_*t*_ (left black axis, open black circles) and stiffness *K*_*t*_ (right grey axis, filled grey squares) reached at the end of the strain phase as a function of *ϕ*_*A*_. The dashed line denotes the predicted scaling for entangled actin networks 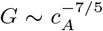. (C) Stiffness *K*(*x*) = *dF/dx* versus *x* (bottom axis) and *t* (top axis) computed from the data shown in (A). The dashed boxed-in region denotes the range of *K*(*x*) values that are averaged to determine *K*_*t*_ values shown in (B). (Inset) Zoom-in of data normalized by the corresponding initial value *K*_0_ for each composite shows the degree to which composites undergo stress stiffening (*dK/dx* > 1) versus softening (*dK/dx* < 1). The horizontal dashed line at *K* = *K*_0_ highlights the degree of stress-stiffening (*K/K*_0_ > 1) versus softening (*K/K*_0_ < 1). (D) Initial force *F*_0_ (left black axis, filled black squares) and stiffness *K*_0_ (right grey axis, open grey circles) measured at the beginning of the strain (*x* = 0) both display non-monotonic dependence on *ϕ*_*A*_. (E) Degree of stiffening *K*_*max*_*/K*_0_ (left black axis, open black circles) and the time over which composites exhibit stiffening 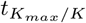 (right grey axis, filled grey squares) increase with increasing *ϕ*_*A*_.

To shed further light on the extent to which elastic contributions to the stress response manifest at different lengthscales, we also evaluate the composite stiffness *K* = *dF/dx* as a function of distance *x* and time *t* = *x/v* (Fig 4C). In the large strain limit (*x* ≳ 2 *μ*m), *K* universally reaches a roughly constant value that increases with increasing *ϕ*_*A*_. We find that this terminal value *K*_*t*_ follows similar power-law scaling as *F*_*t*_ (Fig 4B), corroborating the dominant role that actin plays in persistent stiffness and elasticity of composites. Importantly, this monotonic power-law increase in *F*_*t*_ and *K*_*t*_ with *ϕ*_*A*_ is distinct from the non-monotonic dependence we observe in the linear regime, in which *ϕ*_*A*_ = 0 vimentin networks exhibited stronger elastic signatures than *ϕ* > 0 composites.

We next examined the initial short-time behavior, which we expect to be largely dictated by elastic-like contributions due to the lack of available relaxation mechanisms to appreciably dissipate stress. As shown in the inset of Fig 4A, we observe a dependence on *ϕ*_*A*_ that is quite different than that of both the terminal nonlinear behavior as well as the linear viscoelasticity. Namely, the initial force *F*_0_ is substantially larger for composites compared to single-component networks, in direct opposition to the non-monotonic trend we find for the linear regime metrics *G*^0^ and *η*_0_ which are minimized for composites (Fig 3C,D). We observe similar non-monotonicity for initial stiffness *K*_0_ (Fig 4C,D), corroborating the emergence of enhanced elasticity in composites compared to single-component networks. This emergent behavior may suggest microscale interactions between actin and vimentin [18], depletion-driven restructuring [25], or increased poroelasticity [12, 57–60], ideas we explore further below. Closer examination of *K*(*x*) also reveals modest stress stiffening, i.e., *dK/dx* > 1 for some composites before transitioning to softening *dK/dx <* 0 to a final constant value *K*_*t*_ (Fig 4B). While previous studies have found that vimentin promotes stress-stiffening in cytoskeletal compositess [25, 42–44], we do not find strain stiffening behavior of vimentin networks for the conditions used here, as shown in Fig 4C inset, which shows the *x*-dependent degree of stiffening *K*(*x*)*/K*_0_. The vimentin-only network exhibits no initial stiffening, and the maximum degree of stiffening *K*_*max*_*/K*_0_ as well as the time that the stiffening regime persists, namely the time at which *K*_*max*_*/K*_0_ reaches its maximum, 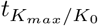, increases monotonically with *ϕ*_*A*_ (Fig 4E). This effect, qualitatively similar to the dependence of *F*_*t*_ and *K*_*t*_, corroborates our understanding that the nonlinear response is dictated primarily by the actin filaments in the composites, which endow composites with stiffness and persistent elasticity for all but the shortest timescales.

### Actin promotes both sustained elasticity and fast relaxation modes in actin-vimentin composites

The nonlinear strain response suggests that actin allows for more sustained elasticity, which we expect to result in reduced relaxation of force following strain as the actin fraction increases. To investigate this conjecture, we measure the time-dependent decay of the force *F* (*t*) that each composite exerts on the trapped probe following cessation of the strain (Fig 5A). We first note that the force at the onset of the relaxation phase, *F* (0), increases with increasing *ϕ*_*A*_ (Fig 5A inset) and displays the same power-law dependence as *F*_*t*_ and *K*_*t*_ (Fig 4B), i.e., 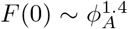 (Fig 4B), as expected as they are measured in close succession, and further supporting the dominant role that actin plays in persistent stiffness.

**FIG. 5.**
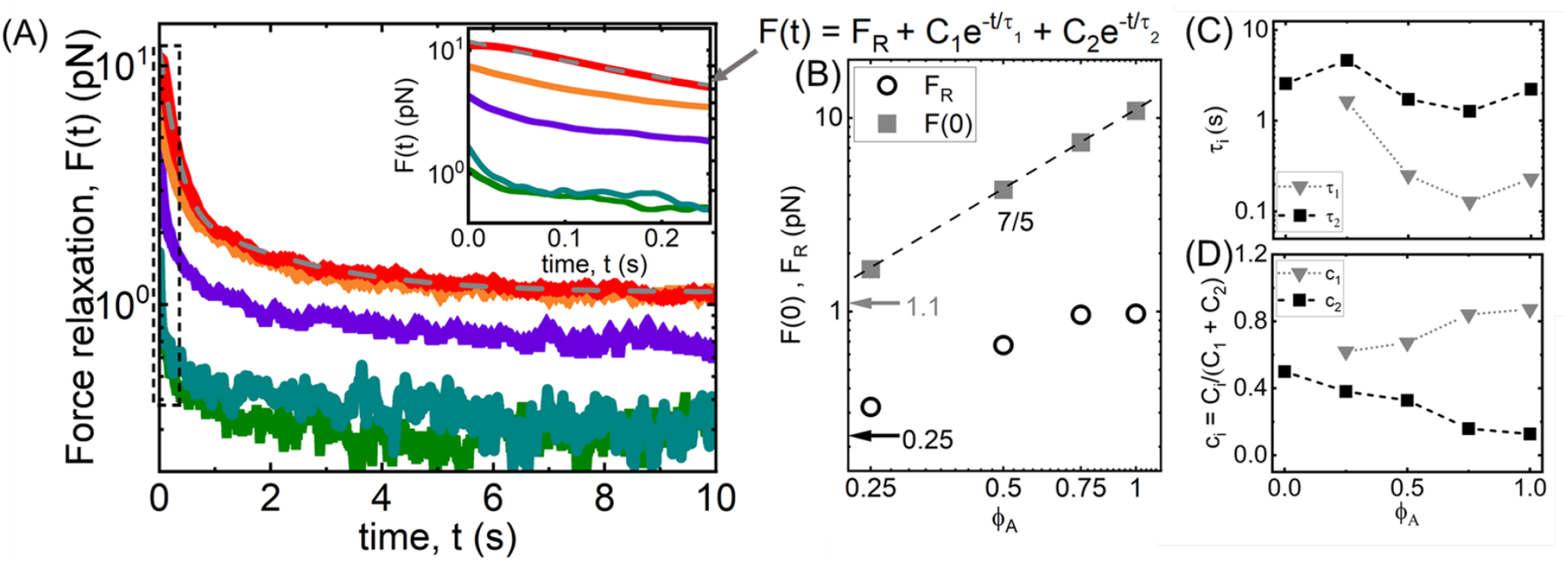
Force relaxation of actin-vimentin composites following nonlinear straining is dictated primarily by entangled actin. (A) Relaxation of force *F* as a function of time *t* following the strain for composites with *ϕ*_*A*_ = 0 (green), 0.25 (dark cyan), 0.5 (purple), 0.75 (orange) and 1 (red). (Inset) Zoom-in of small *t* region denoted by dashed box to show the dependence of the initial force *F*_0_ on *ϕ*_*A*_. (B) Force measured at the beginning (*F* (0), grey squares) and end (*F*_*R*_, black open circles) of the relaxation phase as a function of *ϕ*_*A*_ for *ϕ*_*A*_ > 0 composites. Grey and black arrows and numbers indicate the values of *F* (0) and *F*_*R*_ at *ϕ*_*A*_ = 0. *F* (0) is determined directly from the data and *F*_*R*_ is determined from fitting *F* (*t*) to the function displayed above the plot: 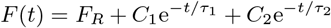. The dashed line represents predicted scaling 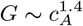 for entangled actin networks. (C) Fast and slow relaxation times, *τ*_1_ (grey triangles) and *τ*_2_, (black squares), as functions of *ϕ*_*A*_, determined from fitting the data in (A) to the function listed in (B). For *ϕ*_*A*_ = 0 data, *τ*_1_ ≈ *τ*_2_ and *C*_1_ ≈ 0, showing that it lacks the fast relaxation that composites with actin exhibit. (D) Fractional coefficients, *c*_*i*_ = *C*_*i*_*/*(*C*_1_ + *C*_2_) associated with the fast (*c*_1_, grey triangles) and slow (*c*_2_, black squares) relaxation modes as a function of *ϕ*_*A*_.

We also evaluate the residual force maintained at the end of the relaxation phase *F*_*R*_, which should equate to *F* (0) and 0 for purely elastic and viscous materials, respectively. Namely, for a purely elastic material, there should be no force relaxation following the strain, such that *F* (*t*) = *F* (0), while a purely viscous fluid would immediately relax all stress once the strain ceases (*F* (*t*) → 0). As shown in Fig 5B, *F*_*R*_ increases with increasing *ϕ*_*A*_, suggestive of increased sustained elasticity, until *ϕ*_*A*_ = 0.75 at which point it saturates at ∼1 pN (∼10% of *F* (0)).

These results demonstrate that actin dictates the ability of composites to exhibit elastic memory following nonlinear straining, which suggests actin is more readily able to withstand large strains, as compared to vimentin, without entanglements being disrupted or filaments being forcibly rearranged. To determine the relaxation mechanisms underlying this behavior, we turn to the time-dependence of *F* (*t*) en route from *F* (0) to *F*_*R*_ (Fig 5A). We find that in the absence of actin (*ϕ*_*A*_ = 0), the force relaxation can be described by a single exponential and offset *F*_*R*_: *F* (*t*) = *Ce*^−*t/τ*^ + *F*_*R*_ with a characteristic decay time of *τ* ≃ 2.6 s. However, all *ϕ*_*A*_ > 0 composites require a sum of two exponentials, 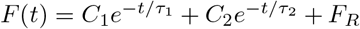, to accurately describe the force relaxation, similar to previous studies on actin networks and composites [12, 35, 61].

The characteristic decay times *τ*_*i*_ and relative coefficients *c*_*i*_ = *C*_*i*_*/*(*C*_1_ + *C*_2_) provide measures of the relaxation times and the degree to which that associated relaxation mechanism contributes to the force relaxation (Fig 5C,D). We first note that the decay time for the vimentin network is comparable to the *τ*_2_ values measured for composites (*τ* ≃ *τ*_2_), so we attribute the decay to the same mechanism. Further, the additional relaxation time that emerges when actin is present, *τ*_1_, is faster than *τ* and generally comparable to 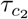 values measured in the linear regime (Fig 3E). The absence of 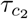 for the vimentin-only network in the linear regime (Fig 3E) is consistent with the lack of *τ*_1_ in the nonlinear regime (Fig 5C), suggesting that it is the same mechanism in both regimes that underlies the fast relaxation and that it is an actin relaxation mode. We also find the *c*_1_ increases with increasing *ϕ*_*A*_, providing further evidence for this phenomenon. As *c*_1_ increases with *ϕ*_*A*_, *c*_2_ by the definition of *c*_*i*_, decreases, reaching < 0.2 for *ϕ* ≥ 0.75. This decrease mirrors the increase in *F*_*R*_, indicating that it is the suppression of the slow relaxation mechanism, likely the disengagement time, that allows for sustained elasticity. Conversely, actin allows for fast relaxation, consistent with the relaxation of filament segments on the order of the confinement tube diameter, which occurs over the entanglement time *τ*_*e*_. More flexible polymers, such as vimentin, are expected to have much faster entanglement times, which are beyond the resolution of our measurement [18]. Alternatively, intermittent crosslinking of vimentin chains would further speed up or altogether eliminate this fast mode. We elaborate on these possible mechanisms below.

## III. DISCUSSION

Our measurements have revealed stark differences between the dependence of composite composition on the mechanical properties of actin-vimentin networks in the linear regime versus the nonlinear regime, as well as at short and long timescales. The nearly opposite emergence of weaker versus stiffer response characteristics of composites as compared to single-component actin or vimentin networks, in the linear versus nonlinear regimes, suggest that different relaxation mechanisms drive the response in the different regimes. More importantly, the non-monotonic dependence highlights the emergent mechanical properties that actin-vimentin composite designs afford that are not a simple sum of the two individual network properties, and may facilitate cellular processes that demand switchable and hierarchical mechanical properties, such as in migration, morphogenesis and division [8–13].

To understand the mechanisms underlying these distinct responses, we first compare the relaxation times that we measure in linear (Fig 3) and nonlinear (Fig 4) regimes (Fig 6A). The linear and nonlinear fast timescales, *τ*_1_ and 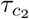, which are absent in the vimentin-only (*ϕ*_*A*_ = 0), are comparable across all *ϕ*_*A*_ values (Fig 6A). As described above, 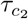 for entangled polymers is a measure of the entanglement time *τ*_*e*_ which, for semiflexible polymers, is predicted to depend on the mesh size *ξ* and persistence length *l*_*p*_ as 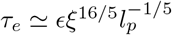 where *ϵ* is the friction term which is expected to be ∼3 s *μ*m^−3^ for actin [33, 62]. We compute the *ϕ*_*A*_-dependent mesh size of the composites *ξ*_*c*_ using the relation 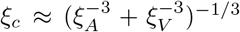 along with 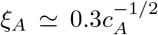 and 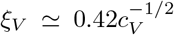 (SI Fig S3) [12, 16]. From these expressions, we compute *τ*_*e*_ values for actin that range from ∼262 ms to ∼120 ms as *ϕ*_*A*_ increases from 0.25 to 1. The *ϕ*_*A*_ = 1 value (*τ*_*e*_ ≃ 120 ms) is comparable to our measured values of *τ*_1_ = 230 ms and 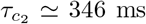, but the increase in these timescales with decreasing *ϕ*_*A*_ is substantially stronger than the predicted dependence assuming a mesh size *ξ*_*c*_. However, if we consider that the actin network appears to dominate the nonlinear response at the end of the strain, we may also expect that the initial relaxation may be likewise dominated by the actin network. As such, if we instead use the mesh size of the actin network *ξ*_*A*_ in the predicted expression for *τ*_*e*_, we compute values of *τ*_*e*_ ≃ 193, ∼369, and ∼1119 ms for *ϕ*_*A*_ = 0.75, 0.5, 0.25, which are comparable to the measured *τ*_1_ values of 129 ms, 249 ms, 1631 ms. The linear regime fast relaxation timescale 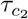 is also of the same order as *τ*_1_ and *τ*_*e*_ for actin but exhibits a weaker and more non-monotonic dependence on *ϕ*_*A*_, resembling the *ϕ*_*A*_-dependence of *ξ*_*c*_ versus *ξ*_*A*_.

**FIG. 6.**
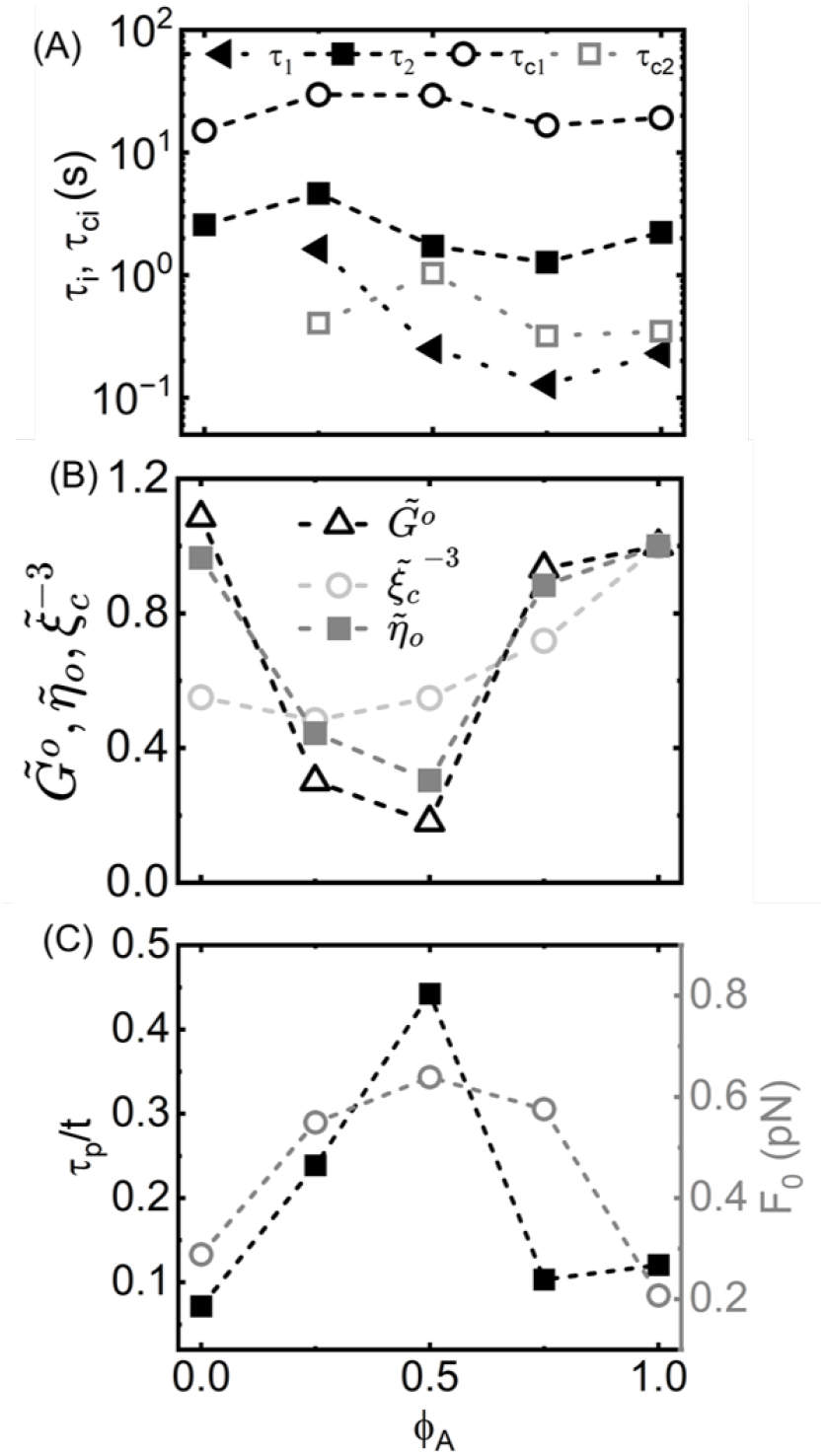
Distinct emergent mechanics of actin-vimentin composites in linear and nonlinear regimes are sculpted by the interplay between actin entanglements, vimentin crosslinking, and poroelasticity. Various metrics and timescales measured in linear and nonlinear regimes versus actin fraction *ϕ*_*A*_. (A) Relaxation timescales from linear (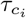, open symbols) and nonlinear (*τ*_*i*_, filled symbols) OTM measurements that correspond to fast (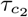, open grey squares; *τ*_1_, black triangles) and slow (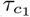, open black circles; *τ*_2_, black squares) relaxation modes. (B) Non-monotonic *ϕ*_*A*_-dependence of measured linear regime metrics, *G*^0^ and *η*_0_, and predicted composite mesh density 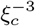, each scaled by their value at 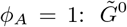 (triangles), 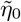 (squares) and 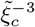 (circles). (C) The ratio of predicted poroelastic relaxation time *τ*_*p*_ to measurement time *t* (left black axis, black squares) and initial force *F*_0_ measured during the nonlinear strain phase (right grey axis, grey circles) both display non-monotonic dependence on *ϕ*_*A*_ with maxima at *ϕ*_*A*_ = 0.5.

Taken together, our results reveal that it is the entanglements between actin filaments that gives rise to the fast relaxation timescale in composites. While the presence of vimentin contributes to the mesh size that dictates the actin relaxation in the linear regime, the relaxation of vimentin itself does not appear to play a role. Moreover, in the nonlinear regime, the relevant confinement network appears to be that of the actin rather than the composite, suggestive of possible forces disentanglement of the vimentin from the actin at the leading edge of the probe [63–65]. This physical picture is consistent with the ostensibly negligible contribution of vimentin to the long-time nonlinear response (Fig 4A) and evidence of forced de-threading in other composite systems [64–66].

Turning to the slow relaxation timescales measured in the linear and nonlinear regimes, 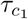 and *τ*_2_, we find that both timescales display similar weak dependence on *ϕ*_*A*_ (Fig 6A). However, *τ*_2_ is generally an order of magnitude faster than 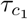. Insofar as we can understand this slow timescale as a measure of the disengagement time, *τ*_*D*_, this faster relaxation in the nonlinear regime is consistent with several previously reported studies of actin networks and composites, in which *τ*_*D*_ was predicted and shown to be faster in the nonlinear regime due to tube dilation and/or entanglement hopping [35, 39, 67]. The former is a result of convective constraint release or forced disentanglement that decreases the local entanglement density and thus speeds up disengagement, while the latter is a process whereby an entangled polymer can hop from one cage to another when an entanglement is transiently disrupted [35, 40, 68]. These mechanisms are direct consequences of nonlinear straining and are largely inaccessible in the quiescent linear regime. The magnitude of *τ*_2_ and its relative insensitivity to concentration is consistent with previous microrheology measurements on actin networks and composites, which were also shown to adopt these non-classical relaxation mechanisms [12, 35]. Further, in the linear regime, *τ*_*D*_ for actin is predicted to be independent of concentration and scale as *τ*_*D*_ ∼ *L*^3^ [33], consistent with the insensitivity of 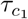 that we uncover.

Given the similar *ϕ*_*A*_-dependence of the relaxation timescales in the linear and nonlinear regimes, coupled with the near opposite dependence of the magnitude of stress response metrics (i.e., *G*^0^, *η*_0_, *F*_0_) on *ϕ*_*A*_ in the two regimes, demands further discussion. In the linear regime, composites appear to be weaker than their single-component counterparts, as summarized in Fig 6B, which shows the plateau modulus and zero-shear viscosity, normalized by their *ϕ*_*A*_ = 1 values, 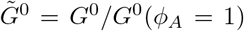 and 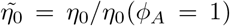. Both quantities follow very similar profiles, reaching minima of ∼0.2 and ∼0.3 at *ϕ*_*A*_ = 0.5. This signature of weaker confinement suggests that the composite mesh size may also exhibit a non-monotonic dependence on *ϕ*_*A*_. Namely, the moduli should scale roughly with the density of the mesh 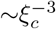 that sets the confinement. Comparing 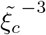 with 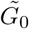 and 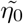 (Fig 6B), we find that 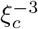 indeed follows a non-monotonic dependence on *ϕ*_*A*_ but its minimum is reached at, *ϕ*_*A*_ = 0.25 rather than *ϕ*_*A*_ = 0.5 and the *ϕ*_*A*_ = 0 value is nearly 2-fold lower than that for *ϕ*_*A*_ = 1 (actin only) and only modestly larger than the minimum at *ϕ*_*A*_ = 0.25. Thus, the decreasing mesh density may explain the drop in moduli values as *ϕ*_*A*_ decreases from 1 to 0.5, but the subsequent rise in values as *ϕ*_*A*_ goes to 0 is substantially more pronounced than the mesh density can account for. We rationalize that the larger than expected value for the vimentin-only network is a result of non-specific crosslinking between VIFs, which has been reported in previous studies [18]. Moreover, previous studies have suggested that introducing actin into vimentin networks may serve to block binding sites and disrupt crosslinking [9, 18], which aligns with the large drop in *G*^0^ and *η*_0_ that we observe between *ϕ*_*A*_ = 0 and 0.25. While previous studies have also indicated potential crosslinking of actin to vimentin, we find no evidence of this interaction, which may be due to the lower filament concentration and ionic strengths we examine compared to this previous study [18]. Crosslinking of vimentin in the absence of actin would also greatly reduce or eliminate *τ*_*e*_, which our data indicate, as crosslinking enhances confinement and reduces the timescale over which filaments can freely diffuse. However, if the crosslinks were long-lived then we may also expect the disappearance of the long relaxation timescale *τ*_*D*_ and terminal flow regime, which we do not observe. This effect indicates that crosslinks are weak and transient, such that the filaments can still exhibit flow-like behavior on long timescales.

The transient and weak nature of the crosslinks likely play an important role in the different responses we observe in the nonlinear regime. The dominant role that actin plays in the nonlinear response (Fig 4A), along with the very weak terminal stiffness *K*_*t*_ and force *F*_*t*_ measured for vimentin networks compared to composites and actin networks (Fig 6B), indicates that the non-linear strain is able to easily disrupt the crosslinks, so they do not contribute to the nonlinear stress response. However, this effect is seemingly at odds with the initial nonlinear response behavior in which composites exhibit more pronounced stiffness and resistance compared to single-component networks, rather than the monotonic increase with increasing *ϕ*_*A*_ that we see for the longer time response features.

To understand these distinct trends, we consider that disruption of vimentin crosslinks is likely to only have a noticeable impact at lengthscales beyond the length between crosslinks, which we can estimate as the vimentin mesh size if we assume that all vimentin crossings contain non-steric linkages. The predicted vimentin mesh size ranges from *ξ*_*V*_ ≃ 0.51 *μ*m to ∼1 *μ*m for *ϕ*_*A*_ = 0 to 0.75. The average vimentin correlation length measured from confocal images is similarly *ξ*_*g,V*_ ≈ 1 *μ*m. Examination of Fig 4A indeed reveals that the distance *x* at which *F* (*x*) for the actin network becomes comparable to and exceeds that of the composites occurs at *x* ≈ 1 *μ*m.

The question remains as to what physical mechanism underlies the emergent stiffness and resistive force *F*_0_ of composites at lengthscales below ∼1 *μ*m (Fig 6C), which is distinct from both the nonlinear response at larger spatiotemporal scales and the linear response. At these very fast timescales (≲100 ms), which are faster than most of the available relaxation modes of the filaments, i.e., *τ*_*e*_, the poroelasticity of the network can contribute to its response to a fast-moving probe[12, 57–60]. Namely, the hydrodynamic drag from a probe of diameter *d* displacing the solvent of viscosity *η*_*s*_ that pervades the network can contribute to the force response at timescales comparable to or lower than the draining time of the fluid, which sets the poroelastic relaxation time 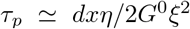. At timescales above *τ*_*p*_ the deformation of the network dominates the response and the hydrodynamic effects of poroelasticity can be neglected. Using our measured *G*^0^ values and computed *ξ*_*c*_ values for each *ϕ*_*A*_, along with *x* = *vt* where *v* = 10 *μ*m s^−1^, we evaluate the ratio of the poroelastic timescale to measurement timescale, 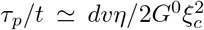, as a function of *ϕ*_*A*_ (Fig 6C). The larger this ratio, the more effect that poroelastic resistance (i.e., slow fluid draining) will have on the initial force response. As shown in Fig 6C, *τ*_*p*_*/t* follows a similar non-monotonic trend as the initial force response *F*_0_, reaching a maximum at *ϕ*_*A*_ = 0.5, suggesting that it is, in fact, the increased poroelastic relaxation timescales of the composites that give rise the emergent increase in initial force response for composites compared to single-component networks. Perhaps counterintuitively, this effect is coupled to the corresponding reduced elastic modulus of composites in the linear regime, demonstrated by the inverse relation between *τ*_*p*_ and *G*^0^. While the mesh size is larger for composites, which lowers the poroelastic ratio via the relation *τ*_*p*_ ∼ *ξ*^−2^, *ξ*_*c*_ drops by less than 30% while *G*^0^ increases ∼6-fold so the increased mesh size has less of an effect.

## IV. CONCLUSION

In conclusion, our suite of optical tweezers microrheology measurements on composites of varying molar fractions of semiflexible actin filaments and flexible vimentin intermediate filaments reveal novel emergent stress response features that are elegantly sculpted by the spatiotemporal scales of the strain. The complex interplay between varying filament stiffness, mesh size, crosslinking, and poroelasticity as the relative molar fractions of actin and vimentin gives rise to seemingly opposite non-monotonic dependence of the response on *ϕ*_*A*_ in linear and nonlinear regimes. This switchable rheology likely plays an important role in cellular processes, such as morphogenesis, adhesion and migration, and may be harnessed for designing multifunctional materials such as soft filtration and sequestration devices. Moreover, our results suggest that transient crosslinking of vimentin dominates the linear viscoelastic response of vimentin-rich composites, while the relative ease with which they can be disrupted hampers the extent to which vimentin fingerprints on the nonlinear response. These weak and switchable non-steric interactions, reminiscent of hydrogen bonding, likely contribute to the broad range and versatility of the mechanical processes that vimentin has been suggested to facilitate in cells.

Notably, our results are at odds with previous bulk rheology measurements that report that vimentin networks exhibit stress-stiffening, owing to their extensibility, and that introducing them into actin networks modulates the response from softening to stiffening by suppressing actin buckling [18, 24]. These differences reinforce the important effect of scale on mechanical properties, in particular for composites with a broad range of characteristic length and time scales. Indeed, the hierarchical mechanical properties of the composite cytoskeleton have been recognized as playing an essential role in its ability to modulate myriad mechanical processes and properties with a finite number of building blocks.

Our future work will explore the effect of ionic strength, which modulates vimentin crosslinking, as well as overall protein concentration, which may tip the scales towards actin-vimentin crosslinking and stiffening behavior in vimentin networks by increasing polymer overlap and entanglement density.

## V. MATERIALS AND METHODS

### Protein preparation

Rabbit skeletal actin (Cytoskeleton, Inc., AKL99) was reconstituted to 46 *μ*M in G-buffer [2 mM Tris pH 8.0, 0.5 mM DTT, 0.1 mM CaCl_2_, 0.2 mM ATP] and stored in single-use aliquots at -80°C. Rhodamine phalloidin (Cytoskeleton, Inc., PHDR1) was reconstituted to 14 *μ*M in methanol and stored at -20°C. Vimentin was prepared by expressing vimentin-coding DNA (pETE7 plasmid provided by Robert Goldman) in Escherichia coli BL21(DE3) component cells, as described previously [69]. Briefly, vimentin was isolated, pooled, dialyzed against a denaturing buffer [50 mM Tris-HCl (pH 8.5), 6 M urea], and flash-frozen for storage at -80 °C. Samples were then dialyzed into a solution containing 5 mM Tris-HCl (pH 8.4) and 1 mM DTT, as described previously [69, 70], by gradually lowering the urea concentration in a step-wise fashion (6M, 4M, 2M, 0M) at room temperature (RT) for 20 min for every dialysis step except the final step which was performed overnight at 4°C.

### Cell culture

Wild-type mouse embryonic fibroblasts (mEFs) were kindly provided by J. Ericsson (Abo Akademi University, Turku, Finland). Cells were maintained in Dulbecco’s Modified Eagle’s Medium (DMEM), including HEPES and sodium pyruvate supplemented with 10% fetal bovine serum (FBS), 1% penicillinstreptomycin, and non-essential amino acids. Cell cultures were maintained at 37°C with 5% CO_2_.

### Immunofluorescence

Cells were fixed for immunofluorescence using 4% paraformaldehyde (Fisher Scientific) for 30 minutes at 37°C. Cell membranes were permeabilized with 0.05% Triton-X (Fischer BioReagents) in PBS for 15 minutes at RT and blocked with 1% bovine serum albumin (BSA) for 1 hour at RT. For vimentin visualization, cells were incubated with primary rabbit anti-vimentin antibody (Abcam) diluted 1:200 in 1% BSA in PBS for 1.5 hours at RT; the secondary antibody anti-rabbit Alexa Fluor 647 (Invitrogen) was used at a dilution of 1:1000 in 1% BSA in PBS for 1 hour at RT. For visualizing actin filaments, cells were stained with rhodamine phalloidin 488 (Invitrogen) diluted 1:200 in 1% BSA in PBS and incubated for 1 hour at RT.

### Actin-vimentin composite preparation

To prepare composites, actin and vimentin filaments were polymerized separately, then mixed and flowed into a microscope sample chamber assembled using double-sided tape as a spacer between a microscope slide and glass coverslip. Actin filaments were polymerized from actin monomers by incubating in an F-buffer [10 mM Imidazole pH 7.0, 50 mM KCl, 1 mM MgCl_2_, 1 mM EGTA, 0.2 mM ATP] at RT for 1 hr. Vimentin IFs were polymerized via the addition of 50 mM Na^+^ and incubation at 37°C for 30 minutes. To form the composite, the solution of polymerized actin was added to the solution of polymerized vimentin, which was then mixed slowly by pipetting up and down ∼10 times before introducing the composite solution into the sample chamber via capillary action. The open ends of the sample chamber were sealed with epoxy and the composites was allowed to equilibrate for 10-15 minutes before collecting data. All composites comprise the same total protein concentration of *c* = *c*_*A*_ + *c*_*V*_ = 11.6 *μ*M with varying molar fractions of actin *ϕ*_*A*_ = *c*_*A*_*/*(*c*_*A*_ + *c*_*V*_) = 0, 0.25, 0.5, 0.75, 1 (Fig 1A). A trace amount of labeled polystyrene microspheres (#FSDG006 dragon green, Bangs Laboratory, Inc.) of 4.2 *μ*m in diameter was added for visualization of the probe particles and microrheology measurements.

### In vitro imaging

To determine the structure of the interpenetrating networks of actin filaments and VIFs in composites, a Leica TCS SP5 laser scanning confocal microscope with 63× 1.4 NA objective was used to image fluorescent-labeled actin and vimentin filaments within composites of the same concentration and *ϕ*_*A*_ values as in microrheology experiments (Fig 1)A. Acti-Stain 555 Phalloidin (Cytoskeleton, Inc) was used to label actin filaments and Alexa Fluor 488 maleimide (Invitrogen) was used to covalently label vimentin, as described in ref. [71]. To reduce photobleaching, oxygen scavenging agents (4.5 *μ*g/ml glucose, 0.005% *β*-mercaptoethanol, 4.3 *μ*g/ml glucose oxidase, 0.7 *μ*g/ml catalase) were included. Stacks of 35 images, each 512 × 512 square-pixels (246.5 × 246.5 *μ*m) in area, with *z* step of 0.5 *μ*m between imaging planes were collected. Each image shown in Fig 2(A) is a mean intensity projection of a representative stack.

To determine structural correlation lengths for the two networks comprising the composites, spatial image autocorrelation (SIA) analysis was performed on each image of each *z*-stack using a custom-written Python script [72– 74]. SIA measures the correlation in intensity *I*(*r*) of two pixels in an image separated by any given radial distance *r* by taking the product of the fast Fourier transform *F* of an image *I* and its complex conjugate, applying an inverse Fourier transform *F*^−1^, and then normalizing by the intensity squared: *g*(*r*) = *F*^−1^(|*F* (*I*(*r*))|^2^)*/*[*I*(*r*)]^2^. The resulting autocorrelation *g*(*r*) generally decays with increasing *r*, and the critical decay distance *ξ*_*g*_ is a measure of the average size of features in the image. For an image of a homogeneous network, *ξ* is often considered analogous to the mesh size. The correlation length *ξ* was determined from our data by fitting the exponentially decaying section of each autocorrelation curve to *g*(*r*) = *g*_*o*_ exp(−*r/ξ*), as described previously [75, 76].

### In vivo Imaging

Vimentin and actin images were obtained using spinning disk confocal microscopy (Yokogawa CSU-W1) on an inverted Nikon Ti-E microscope and 100× oil immersion objective (1.49 NA) imaged onto an Andor Zyla CMOS camera.

### Optical Tweezers Microrheology

All optical tweezers microrheology (OTM) measurements were performed using an optical trap built around an IX73 fluorescence microscope (Olympus, Melville, NY) with a 1064 nm Nd: YAG fiber laser (BKtel, RPMC Lasers, Inc) focused with a 60× 1.42 NA oil immersion objective lens (UPLXAPO60XO 60x NA 1.42, Olympus). The force was measured using a position-sensing detector (PSM2-10Q PSD module coupled with amplifier OT-301, ON-TRAK Photonics, Inc) that recorded the deflection of the trapping laser, which is proportional to the force acting on the trapped probe. The trap stiffness, which relates the laser deflection to force, was measured using Stokes drag [77] and passive equipartition calibration methods [78, 79].

We performed average linear OTM measurements of 20 trials by 20 beads at 20 different locations in the sample by recording the thermal fluctuations of an optically trapped probe for 135 seconds at a frequency of 20 kHz. From this data, we extracted the storage modulus (*G*^′^(*ω*)) and loss modulus (*G*^′′^(*ω*)) as a function of the angular frequency *ω* via the generalized Stokes-Einstein relation (GSER) as described in reference [80]. In brief, the normalized mean-squared displacement of the ensemble of trapped beads, 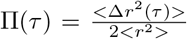, was computed, after which 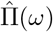, the Fourier transform of Π(*τ*), was obtained via the relationship explained in Tassieri et al.[80]:

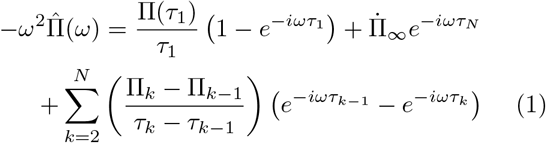

where 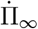 is the slope of Π(*τ*) extrapolated to infinite time, *τ* is the lag time, *τ*_*N*_ is the *N* ^*th*^ lag time, and 1 and N represent the first and last point of the oversampled Π(*τ*). The PCHIP function in MATLAB was used to perform oversampling. The complex modulus, *G*^*^(*ω*) = *G*^′^(*ω*) + *iG*^′′^(*ω*), was then determined from 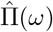 as

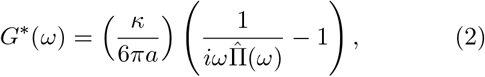

where *a* is the bead radius and *κ* is the trap stiffness. From *G*^′^ and *G*^′′^, the complex viscosity *η* = (*G*^′2^ + *G*^′′2^)^1*/*2^*/ω*, and loss tangent tan *δ* = *G*^′′^*/G*^′^ were also computed. All data shown in Fig 3 are averages over 20 trials performed with different beads in different regions of the sample.

To perform nonlinear OTM measurements, an optically trapped probe was displaced a distance of 10 *μ*m through the composite at a constant speed of *v* = 10 *μ*m s^−1^ using a nanopositioning piezoelectric stage (PDQ-250, Mad City Laboratories) to move the sample chamber relative to the trap. Following this 1 s strain phase, the trap is held fixed for 15 s while the probe relaxes back towards the center of the trap. The laser deflection is recorded at 20 kHz during both the strain and relaxation phases (Fig 1B,D). The force curves displayed in Figures 4 and 5 are averages over 30 trials using 30 different probes at different locations in the sample chamber. The actin concentration and strain rate were both chosen to be higher than the previously determined concentration and strain rate necessary for the onset of nonlinear mechanics [37]. Custom LabVIEW (National Instruments) and Matlab codes were used for all instrumentation control, data acquisition, and data analysis.

## Supporting information

https://drive.google.com/file/d/1I6CQlCfflNZ3LBJaxs1Dk23xahnfMgDb/view?usp=drive_link

## ACKNOWLEDGMENTS

We thank Bobby Carroll for supplying vimentin samples and Karthik Peddireddy for assistance with OTM acquisition and analysis software. This research was funded by Bucknell University (BJG), NSF DMREF 2119663 and NIH R15GM123420 to RMRA, and NSF CMMI 2238600 and NIH R35GM142963 to AP.

JP and RS performed experiments; AP provided guidance on experiments and materials, helped interpret data, and reviewed and edited the manuscript; RMRA provided guidance on experiments and analysis, interpreted data, and wrote the manuscript; BJG designed and guided experiments, analyzed and interpreted data, and wrote the manuscript.

The authors declare no conflict of interest.

