## Supplementary material for "Switchable microscale stress response of actin-vimentin composites emerges from scale-dependent interactions": https://drive.google.com/file/d/1I6CQlCfflNZ3LBJaxs1Dk23xahnfMgDb/view?usp=drive_link

(Dated: June 6, 2024)

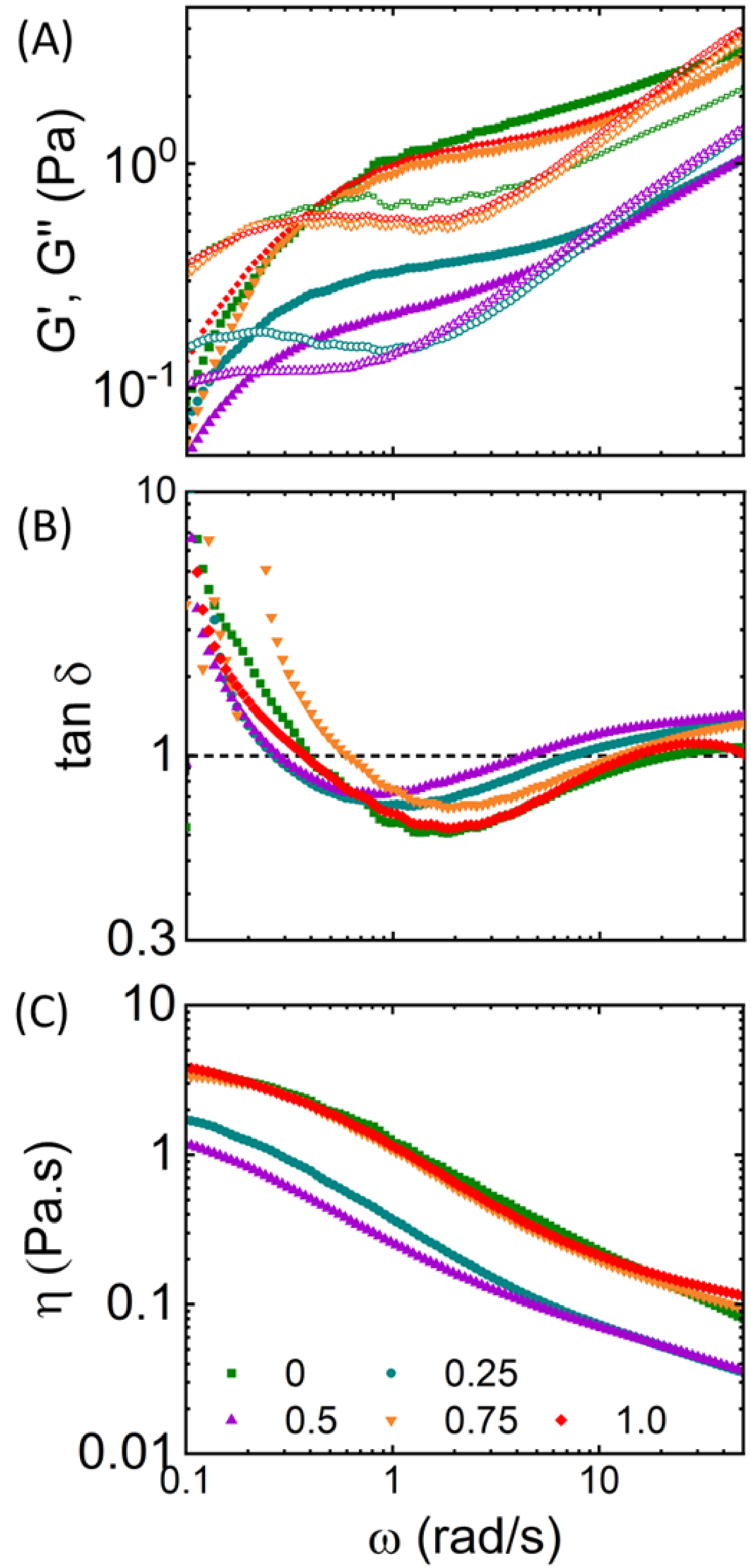


FIG. 1. S1. (A) Linear frequency-dependent elastic modulus $G'$ (closed symbols) and viscous modulus $G''$ (open symbols) of the actin-vimentin composite networks, (B) Frequency-dependent loss tangent $tan\ \delta = G''(\omega)/G'(\omega)$ versus $\omega$ computed from data shown in (A) with a dashed black line representing $G' = G''$, indicating the extent to which each system exhibits elastic versus viscous dynamics of F-actin-vimentin composite, (C) Frequency-dependent shear viscosity $\eta(\omega)$. Each composition is the average of 20 different trials, as explained in the methods section of the main text.

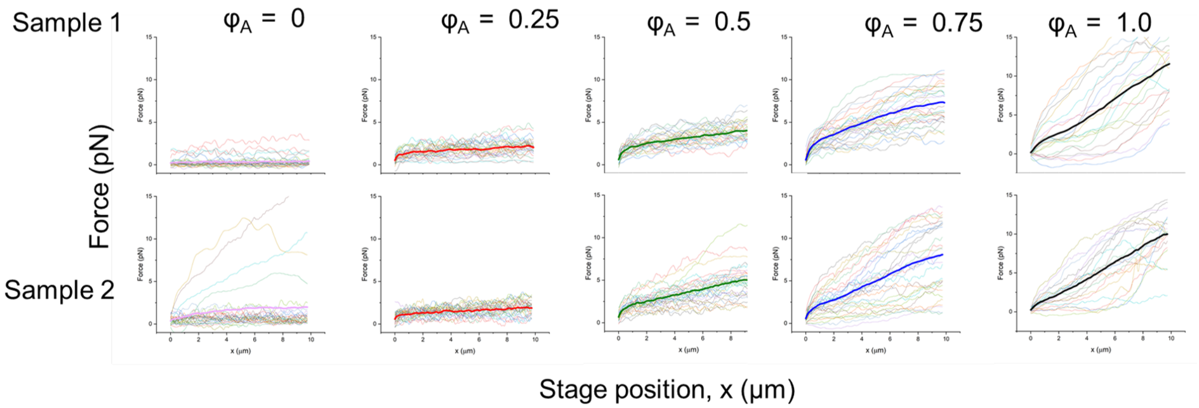


FIG. 2. S2. Individual force trials that are averaged over to compute the average force curves F(x) shown in Fig 4A. Each trial uses a different microsphere probe in a different region of the sample chamber. Each panel corresponds to a different combination of $\phi_A$ (columns) and the two replicates (rows) as indicated on the top and left of the panel grid.

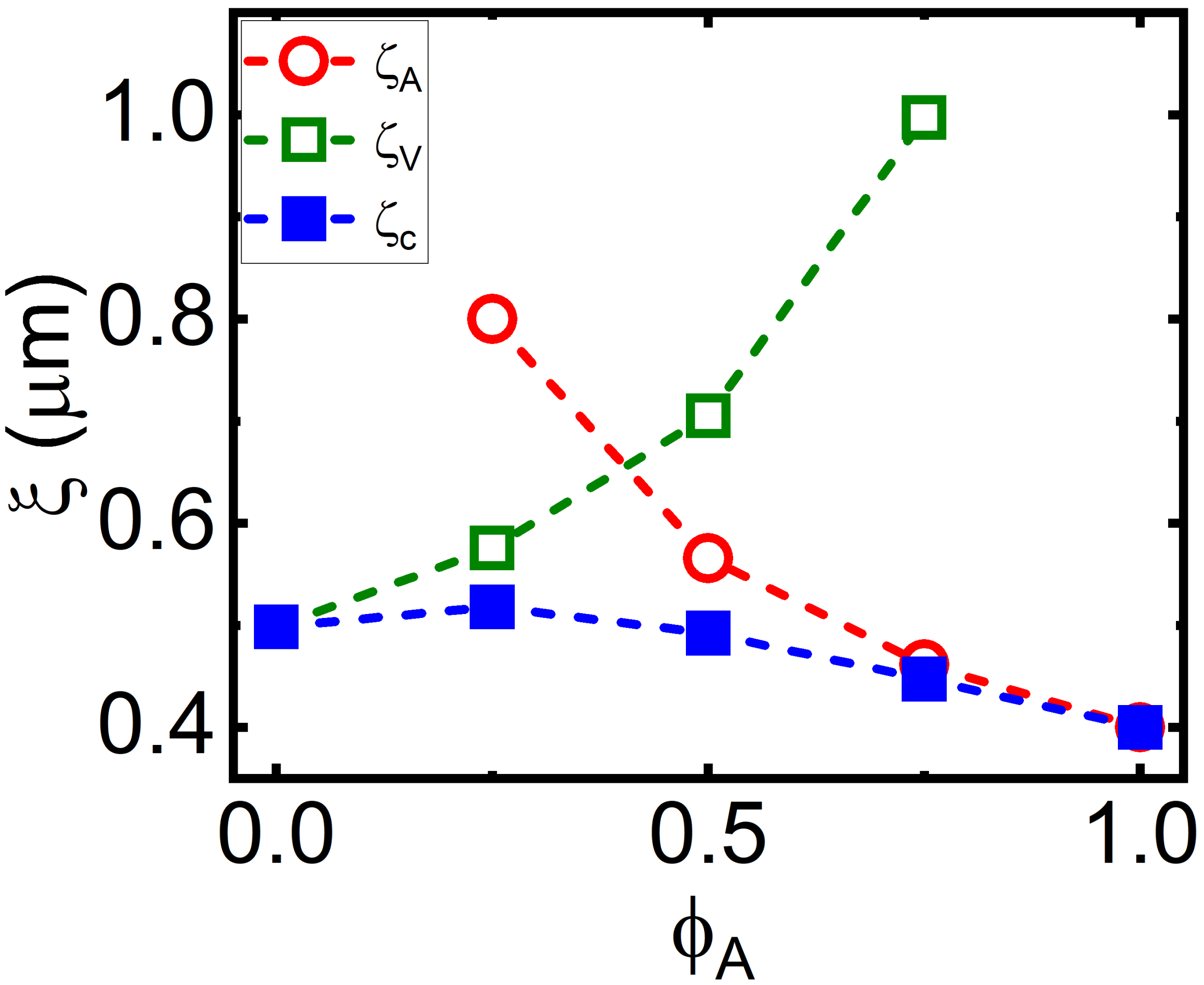


FIG. 3. S3: The computed mesh sizes of actin ($\xi_A$), vimentin ($\xi_V$), and the actin-vimentin composite ($\xi_c$) networks, which represent the average distance between adjacent entanglement points, were determined as described in the main text. The mesh size of actin, $\xi_A$, decreases as the concentration of actin ($\phi_A$) increases, while the mesh size of vimentin, $\xi_V$, increases. However, the overall mesh size of the composite, $\xi_c$, remains approximately the same ($\xi_c = [\xi_A^{-3} + \xi_V^{-3}]^{-1/3} \approx 0.5$ $\mu$m).
